# Metagenomic analysis uncovers strong relationship between periodontal pathogens and vascular dysfunction in American Indian population

**DOI:** 10.1101/250324

**Authors:** Prathik K Vijay Kumar, Roberta A. Gottlieb, Suzanne Lindsay, Nicole Delange, Tanya E. Penn, Dan Calac, Scott T. Kelley

## Abstract

Periodontal disease (PD) is a well-known risk factor for cardiovascular disease (CVD) but the casual relationship is unclear. American Indians/Alaskan Natives (AI/AN) have high rate of both PD and CVD and a better understanding of how PD might affect heart health would be particularly helpful in this population. In this study, we sequenced the bacterial biofilms of periodontal (gum) pockets and used metagenomic sequencing and vascular health measurements (immune cytokine profiles and vascular flow) to determine the relationship of microbial pathogens and CVD. Twelve subjects were sequenced before and after standard periodontal treatment. Other measures taken before and after treatment included a full dental screening; serum concentration of key immune cytokines from blood samples; lipid profiles from fasting venous blood; and plasma glucose concentrations. The non-invasive Laser Doppler Fluxmetry (LDF) procedure was conducted to measures the microvascular vasodilation. We found highly significant relationships between the total abundance of 4 periodontal pathogens, *Porphyromonas gingivalis, Fusobacterium nucleatum, Tannerella forsythia and Treponema denticola*, and the inflammatory cytokine interleukin 1 beta (IL-1β) (r=0.63; p=0.009) as well as with vascular flow post sodium nitroprusside (SNP) treatment (r=p=0.006). Two bacterial species that correlated most with IL-1β were *F. nucleatum and P. gingivalis*. IL-1β has been strongly implicated as a causal factor in atherosclerosis and in periodontal bone loss. To our knowledge, this is the first direct link between abundance of specific periodontal pathogens and cardiovascular disease in humans, and suggests that these pathogens could be used as warning signs for cardiovascular risk.

## Introduction

Periodontitis (gum disease) is a polymicrobial, chronic condition inflammatory disease of the gingiva characterized by a progressive breakdown of gum tissue. If left untreated, periodontitis results in a deterioration of the alveolar bone resulting in loosening and eventual loss of the tooth (1,2). Oral microbiologists have determined that periodontal disease (PD) is caused by complex biofilms that form at the tooth-gum interface. As the disease progresses, the biofilms become more complex with the early colonizers being mostly commensal bacteria (e.g., *Streptococcus* and *Veillonella*), and the later colonizers tending to be more pathogenic (e.g., *Porphyromonas* and *Treponema*) (2-4).

Numerous studies have established risk factors for coronary artery disease including smoking, obesity, diabetes, hypertension and hypercholesterolemia (5). In addition, a number of studies over the past two decades have suggested an association between periodontitis infection and cardiovascular disease (CVD) (6-9). A meta-analysis found a 44% increase in risk of future cardiovascular disease in adults under age 65 with periodontal disease (10), while another study found a clear association between chronic periodontitis and risk of coronary heart disease that revealed a hazard ratio of 2, independent of all other cardiovascular risk factors (6). Another study of endothelial function, a reliable indicator of vascular health, and serum inflammatory markers in middle-aged men showed improvement of vascular function after aggressive treatment of their periodontal disease (11). A 2007 meta-analysis by Bahekar *et al.* of cohort studies, case-control studies and cross-sectional studies, found that individuals with PD had between a 1.14 and 2.2 times greater risk of developing coronary heart disease (CHD) than those without PD (12).

PD is hypothesized to contribute to atherosclerosis in two ways. First, the pathogens causing periodontal disease may directly infect the atherosclerotic plaques. Direct studies of atherosclerotic plaques have detected the presence of *Porphyromonas gingivalis*, *Streptococcus sanguis*, and other major oral bacteria (13,14). However, this may represent secondary colonization of pre-existing atherosclerotic plaques at sites of turbulent flow. Some studies have also raised the possibility that bacterial invasion of the endothelium occurs and several studies have documented periodontal pathogens present in atherosclerotic plaques (15,16).

Second, periodontitis can lead to systemic increases in inflammatory and immune responses that can indirectly contribute to atherosclerosis (17-19). The presence of periodontitis is accompanied by a local inflammatory response, with invasion of neutrophils and lymphocytes. However, because of its chronicity, this is thought to progress to systemic inflammation, and it has been shown that patients with periodontitis have elevated levels of tumor necrosis factor-alpha (TNFα), interleukin 1 beta (IL-1β), C-reactive protein (CRP), interleukin 6 (IL-6), and monocyte chemoattractant protein 1 (MCP-1/CCL2) (17-19). A 17.5 years follow-up study of 2549 individuals with major coronary heart disease and 3696 control subjects evaluated whether CRP and other inflammatory markers could serve as biomarkers for CHD risk. The study concluded that CRP is a moderate predictor for CHD (20). It is not clear whether the increase in inflammatory cytokines represents a response to chronic local infection or to the presence of bacteria (or bacterial fragments) circulating in the blood, but in either case, a systemic inflammatory response is initiated.

Atherosclerosis itself is increasingly being regarded as an inflammatory disease, and patients with other systemic inflammatory disorders such as systemic lupus erythematosus or rheumatoid arthritis have accelerated atherosclerotic disease (21-22). It seems reasonable to conclude that the systemic inflammation arising from periodontal disease would contribute to the development or progression of atherosclerosis. Magnitude of the inflammatory burden has also been shown to be related to the progression of atherosclerotic calcifications in patients with advanced disease (12,23). A study in mice by Chukkapalli *et al.* (2015) found experimental evidence connecting specific PD pathogens with both systemic inflammatory processes associated with CVD and evidence of aortic bacterial inflammation (24). The researchers infected the oral cavity of ApoE^Null^ mice with a bacterial consortium of well-established human periodontal pathogens: *Porphyromonas gingivalis, Treponema denticola, Tannerella forsythia* and *Fusobacterium nucleatum*. An analysis of the mice over a 24-week period determined clear evidence of PD as well as increases in serum risk factors associated with atherosclerosis, a reduction in serum NO which is indicative of endothelial dysfunction, elevated serum cytokine levels, enhanced aortic plaque development, and even evidence of live bacteria inside aortic plaques (24). The researchers also reported that the combination of pathogens elicited a much stronger immune response than previous studies with single pathogens, suggesting a synergistic effect of the polymicrobial consortium.

In this study, we used direct DNA extraction and metagenomic sequencing to explore the relationship of periodontal biofilm bacteria to levels of systemic inflammation and vascular dysfunction in an American Indian/Alaska Native (AI/AN) population in southern California. AI/AN populations have a higher prevalence of both periodontal disease and heart disease than the general population, making the potential connection between gum disease and heart disease an important subject of study for this population. This study builds upon our previous amplicon-based 16S ribosomal RNA marker gene study, which focused solely on periodontal disease. While single-marker gene analysis is highly useful for determining species composition in microbial communities, shotgun metagenomic analysis of whole microbial communities provides more refined species and strain identification as well as insight into the functional gene composition of biofilm communities. Our results showed that the abundances of four primary periodontal pathogens associated with atherosclerosis correlated with the blood serum immune system cytokine profiles and vascular function measurements from the same participants. Our analysis also uncovered strong relationships between the relative abundance of PD pathogens and immune factors previously associated with atherosclerosis and periodontal disease, as well as Laser Doppler fluxmetry (LDF) measurements of vascular function.

## Materials and methods

### Ethics Statement

This study was conducted as a partnership by San Diego State University (SDSU) Institute of Public Health (IPH) and SDSU Bioscience Center. The study was approved by the SDSU and Southern California American Indian Health Center institutional review boards, and participants were enrolled after informed consent was obtained. Written informed consent was obtained from each participant. The study was registered as “Assess the Effect of Treating Periodontal Disease on Cardiovascular Function in Young Adults” on ClinicalTrials.gov under the identifier NCT01376791.

### Study Population

American Indian/Alaska Native (AI/AN) adults age 21-40 seeking medical or dental care at a clinic in southern California were invited to participate in the study. Participants with known preexisting heart disease conditions (hypertension, atherosclerosis, valve disease, peripheral vascular disease, history of acute myocardial infarction, or heart failure), known inflammatory conditions (autoimmune disorders, chronic infections outside the mouth) and immunosuppression (due to medication or infected HIV) were all excluded. Other exclusion criteria included: recent periodontal treatment, antibiotic use in previous three months, high blood pressure (> 90mm Hg), ongoing use of anti-inflammatory medication (other than H2 blocker), statins or medications that affect periodontal status (phenytoin, calcium antagonists, cyclosporin, coumarin, heparin), and or dentition with fewer than 20 teeth. After careful assessment of the physical and dental health of the participant and consideration of the inclusion and exclusion criteria, 36 total participants with periodontitis were available for the study. Of the original 36 participants, we selected 24 total samples from 12 participants, 1 pre‐ and 1 post-periodontal treatment (12 participants X 2 samples = 24 total samples) to subject to shotgun metagenomic sequencing. The selection of the 12 samples was based on having an equal number of individuals having a clear response to periodontal treatment as measured by significant changes in gum pocket depth post-treatment: 6 patients improved after treatment (average periodontal pocket depth decreased) and 6 patients worsened after treatment (average periodontal pocket depth increased).

### Clinical Examination

Two specially trained registered dental hygienists assessed periodontal disease status was using standardized method. Based on the periodontal pocket depth (PPD), clinical attachment loss (CAL), plaque score, and bleeding on probing (BOP), participants were classified to various degrees of periodontitis. Patients with PPD≤4 mm, CAL≤3 and BOP>10% were classified as having gingivitis. With PPD≥5 mm, CAL≥4 and BOP≥30%, patients were classified as having mild-moderate periodontitis, and patient with PPD≥7 mm, CAL≥6 and BOP≥30%, were classified as having severe periodontitis. Height and weight were obtained with the subject lightly clothed and without shoes. Body mass index was calculated as the ratio of weight in kilograms divided by height squared in meters (kg/m^2^). Information about medical history, health behaviors and demographics were obtained from the participants using a questionnaire that was administered during an interview by research staff.

### Laboratory Analysis

Blood samples were obtained by a nurse one week after enrollment to determine serum concentration of CRP, IL-6, interleukin 10 (IL-10), IL-1ß, TNF-α, Interferon-γ (IFN) and Cardiac Troponin I (cTnI). Blood was also collected to determine high-density lipoprotein (HDL), low-density lipoprotein (LDL) and total cholesterol levels. Samples were processed immediately and stored at −70°C until analyzed by commercial laboratory. Singulex Clinical Laboratory (Alameda, CA) used a Roche analyzer for CRP measurements. cTnI, IL-6, IL-10, IL-1ß, IFNɣ and TNF-α were analyzed using SMC Erenna Immunoassay with single-molecule counting technology on kits 03-0092, 03-0089, 03-0056, 03-0028, 03-0049 & 03-0088 respectively. Non-invasive LDF measures endothelial function reflected by microvascular vasodilation; endothelial dysfunction is considered an early predictor of atherosclerosis. Life Tech iontophoresis device was used to infuse Acetylcholine (Ach) and Sodium nitroprusside (SNP) for 10 minutes, and the microvascular vasodilation were recorded at 1, 5 and 10 minutes. Area under the curve for these LDF-recordings (SNP10 and Ach10) was integrated and expressed calibrated Amplitude Units x time, defined as the total number of blood cells passing two points 10 minutes after SNP treatment. The results from these instruments were printed, images and tracings were analyzed by technicians using appropriate software, and LDF integrated values are documented. All studies were repeated 3 months after the participants completed periodontal treatment.

### Metagenomic Sequencing and Bioinformatics

PCR samples of the 12 subjects before and after treatment (total of 24 PCR samples) were submitted to the core facility at The Scripps Research Institute (TSRI) for metagenome sequencing on Illumina MiSeq. Fastq sequence files were converted to fasta format and were uploaded to MG-RAST. After performing the default sequence quality control using MG-RAST and *removing any detectable human sequence contamination*, taxonomic classification, organismal abundance and functional abundance were assigned to sequences by MG-RAST.

Raw sequences, MG-RAST output and study metadata are available on MG-RAST, ID number MGP15104 (http://metagenomics.anl.gov/linkin.cgi?project=mgp15104). Relative abundance of *Porphyromonas gingivalis*, *Fusobacterium nucleatum*, *Tannerella forsythia*, and *Treponema denticola* were calculated by dividing the abundance counts of individual sequences determined to belong to these four species by the total sequences in each metagenomic sample. For the statistical analyses, we used both the relative abundance of each individual pathogen and also calculated the “total abundance” for all four pathogens, which was the sum of the relative abundance of all four periodontal pathogens.

### Statistical Analysis

Frequencies for all categorical variables are reported. Means and standard deviations are presented for continuous variables, and the median and interquartile ranges are reported for non-normally distributed data. Cytokines and vascular functions values had skewed distributions and were log-transformed to normalize the data. Pearson’s product-moment correlations were used to determine if relative pathogen abundance correlated with cytokine, lipid and LDF measurements. Bivariate association between total relative abundance of the periodontal pathogens and each of the cytokines and vascular functions were examined using simple linear regression with 95 percent confidence. Bonferroni corrections were used to adjust for multiple comparisons. Statistical analyses were performed using R (version 3.3.2) in a Jupyter notebook web interface (version 4.2.1). Log2fold analysis of MG-RAST data was performed using RStudio (version 9.4) with libraries from Bioconductor (version 3.4). The data files, Jupyter notebooks, and the R command files are included as supplemental files (S1 File).

## Results and discussion

Table 1 details the general characteristics of the population whose periodontal biofilms were analyzed in this study. Our metagenomic analysis of serum cytokines and periodontal pathogens uncovered a statistically significant relationship between the relative abundance of four primary periodontal pathogens and IL-1β (Pearson correlation; r=0.64, p=0.0008, p(adj)=0.006; Table 2). We also discovered a positive relationship with IFNγ, but the association was not significant post correction (r=0.42, p=0.039, p(adj)=0.27; Table 2). These findings are somewhat in contrast to the correlations between cytokine levels and measure of PPD in a parallel study of the same population. Delange et al. (*in review*) found an association between PPD and the blood level cytokine concentration of CRP and IL-6 although these were not significant after adjusting for possible confounders. The strong correlation between pathogenic members of the periodontal biofilm and IL-1β cytokine levels suggests that a more targeted focus on specific members of periodontal pocket biofilms and their relationship to the immune response could more directly connect periodontal disease and heart health.

**Table 1.** Population characteristics.

| Characteristics | n(%)<sup>a</sup> |
| --- | --- |
| **Periodontal Status** |  |
| Severe | 4 (33.3%) |
| Moderate | 5 (41.7%) |
| None/Mild | 3 (25.0 %) |
| **Gender** |  |
| Male | 6 (50%) |
| Female | 6 (50%) |
| **Characteristics** | **Mean(SD)** |
| Age, yrs | 28.8 (6.9) |
| BMI<sup>b</sup>, kg/m<sup>2</sup> | 30.7 (7.5) |
| **Cytokines** | **Median(SD)** |
| CRP (mg/L) | 3.7 (9.4) |
| cTnI (pg/mL) | 0.6 (0.9) |
| TNFα (pg/mL) | 2.0 (0.9) |
| IL-10 (pg/mL) | 1.8 (0.6) |
| IL-6 (pg/mL) | 1.4 (1.0) |
| IFNγ (pg/mL) | 0.2 (0.2) |
| IL-1β (pg/mL) | 0.2 (0.2) |
| **Lipid Profile** | **Mean (SD)** |
| HDL | 193.7 (43.8) |
| LDL | 46.5 (13.8) |
| CHOL | 109.3 (39.5) |
| TRI | 164.2 (105.9) |
|  | **Median (IQR)** |
| SNP 10 | 43309.7 (30059.3) |
| ACh 10 | 72965.7 (85331.8) |
|  | **Mean(SD)** |
| SNP 10 | 24983.1 (11910.7) |
| ACh 10 | 95053.9 (67601.9) |

General characteristics of the population subset analyzed in this study.
<sup>a</sup> Results from 6 Males, 6 Females were combined pre- and post- treatment for Means, Medians and SDs.
<sup>b</sup> BMI=Body Mass Index.

**Table 2.** Correlations between periodontal pathogens and cytokine levels.

| Biomarker | r | p-value | p-value (adj)<sup>a</sup> |
| --- | --- | --- | --- |
| CRP | 0.03 | 0.88 | 1.0 |
| cTnI | 0.37 | 0.07 | 0.84 |
| TNFα | 0.11 | 0.61 | 1.0 |
| IL-10 | 0.08 | 0.71 | 1.0 |
| IL-6 | 0.09 | 0.69 | 1.0 |
| IL-1β | 0.64 | 0.0008** | 0.006** |
| IFNγ | 0.42 | 0.039* | 0.27 |

Pearson's product-moment correlations of total relative abundance of four periodontal pathogens (ln-transformed) with cytokines measurements (ln-transformed).
<sup>a</sup> P-value (adj) Bonferroni corrected for multiple comparisons.
\* p < .05, \*\*p < .01.

The positive correlation between the periodontal pathogens and high IL-1ß is consistent with the apparent role of this cytokine in PD. IL-1ß is known to play a critical role in stimulating bone resorption in the later stages of periodontal disease (25,26) (bone loss through resorption happens in the final stages of severe periodontitis) and a study of adults with periodontal disease found higher average concentration of IL-1ß in patients with severe periodontal disease than in those with a healthy periodontium (27). Furthermore, the connection between periodontal disease and CVD, particularly atherosclerosis, via serum IL-1ß appears to have significant support in the literature. Several previous studies have strongly associated IL-1β levels with cardiac health, and even indicated a mechanistic link in animal models. A 2003 population-based study of 1,292 subjects, found IL-1ß level to be four times higher in subjects diagnosed with congestive health failure (28). A study in mice also evaluated the effect of IL-1ß on formation of atherosclerosis and determined that IL-1ß promoted atherosclerosis (29). Other animal studies have shown that deleting or inhibiting IL-1 signaling (either by administration of exogenous IL-1ß or by blocking IL-1Ra) reduced formation and progression of atherosclerotic plaques, strongly indicating a mechanistic link (30,31).

Bivariate analysis determined that, of the four pathogens, the relative abundance of *F. nucleatum* had the strongest relationship to blood serum IL-1ß levels (Table 3). This is congruent with previous research on periodontal disease which specifically identified *F. nucleatum* as a strong stimulator of neutrophil secretion of IL-1ß (32,33). Our results, in combination with the animal model studies and the periodontal research, indicate this organism contributes to both periodontal disease and inflammatory processes associated with vascular dysfunction.

**Table 3.** Associations of specific periodontal pathogens and IL-1β.

| Abundance | Coefficient | Std. Error | t | P- Value (adj)<sup>a</sup> |
| --- | --- | --- | --- | --- |
| *Porphyromonas gingivalis* | 0.25 | 0.22 | 1.12 | 0.28 |
| *Tannerella forsythia* | 0.04 | 0.18 | 0.21 | 0.84 |
| *Treponema denticola* | -0.02 | 0.12 | -0.19 | 0.85 |
| *Fusobacterium nucleatum* | 0.33 | 0.16 | 2.0 | 0.05 |

Bivariate Analysis of relative abundances of individual pathogens (ln-transformed) with IL-1β measurements (ln-transformed).
<sup>a</sup> P-value (adj) Bonferroni corrected for multiple comparisons.

A weak correlation was detected between blood cholesterol (CHOL) levels and total relative periodontal pathogen abundance, but the adjusted p-value was not significant (Table 4). No significant correlations were found with other blood lipid profiles. These results agree with a case-control study which evaluated the relationship between periodontitis and serum lipid profile in 60 patients, 30 with and 30 without chronic periodontitis (34). Blood serum levels of total cholesterol, triglycerides (TRI), HDL and LDL were measured in both groups and there was no significant difference in any measure of blood serum lipid concentration.

**Table 4.** Correlations between periodontal pathogens with lipid profile measurements.

| Lipid Profile | r | P-value | P- Value (adj)<sup>a</sup> |
| --- | --- | --- | --- |
| CHOL | 0.44 | 0.03* | 0.39 |
| HDL | 0.08 | 0.71 | 1.0 |
| LDL | 0.29 | 0.79 | 1.0 |

Pearson's product-moment correlation of total relative abundance of four periodontal pathogens (ln-transformed) with lipid profile measurements (ln-transformed).
<sup>a</sup> P-value (adj) Bonferroni corrected for multiple comparisons.
\* p < .05.

Table 5 shows the relationship between direct LDF vascular function measurements and total relative abundance of periodontal pathogens. A strong and significant relationship was found between total relative pathogen abundance and LDF measures of blood flow volume after 10 minutes of Sodium Nitroprusside (SNP) treatment (p=0.0005). The SNP10 value is an estimate of the total number of blood cells passing between two points 10 minutes after SNP treatment. The values after 1 minute and 5 minutes were also negatively correlated (data not shown). Interestingly, there was no correlation with Acetylcholine (Ach) treatment. We are not clear why this did not mirror the SNP result, though we note that Ach stimulates the release of endogenous nitric oxide, while SNP is based on the addition of an exogenous nitric oxide source. Although any given measure of vascular function is not without controversy, our data are consistent with a relationship of relative periodontal pathogen load and poorer vascular function.

**Table 5.** Correlations between periodontal pathogens with LDF measurements.

| LDF measurements | r | P-value | P-value (adj)<sup>a</sup> |
| --- | --- | --- | --- |
| SNP10 | -0.65 | 0.0005** | 0.0065** |
| Ach10 | 0.06 | 0.79 | 1.0 |

Pearson's product-moment correlation for total relative abundance of four periodontal pathogens (ln-transformed) with LDF measurements.
<sup>a</sup> P-value (adj) Bonferroni corrected for multiple comparisons.
\*\* $p < .01$ .

Metagenomics analysis also allows the opportunity to investigate the gene functional diversity of microbial communities per se, and the relationship of gene functional pathways to inflammation or disease. Given the strong association of total relative pathogen abundance to IL-1ß levels, we asked whether the abundances of any given gene pathway differed significantly between the periodontal pocket microbiomes of individuals with high IL-1ß levels and those with low IL-1ß levels. Fig 1 shows the results of a fold change (log2) analysis based on estimated relative abundances of functional gene pathways. Over 40 different function categories differed between the microbiomes of individuals with high and low IL-1ß. The bacteria in individuals with low IL-1ß tended to be more abundant in genes involved in metabolism and biosynthesis. The bacterial communities in high IL-1ß individuals, on the other hand, were richer in degradation (Histidine) and cell death pathways.

**Fig 1.**
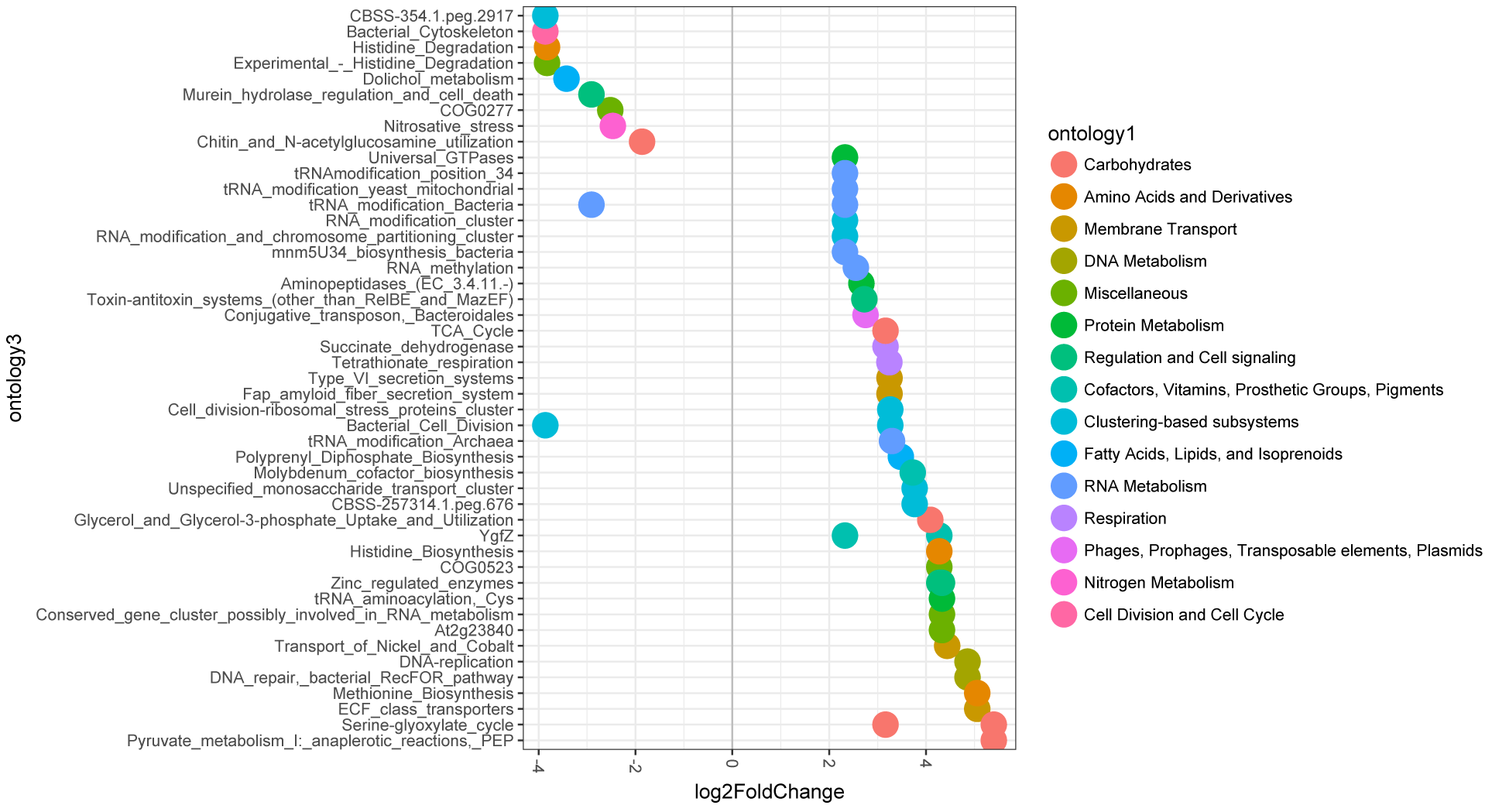
Plot showing log2 fold-change values (x-axis) by gene functional categories. The relative abundance is significantly different (adjusted *P*<0.05, >two-fold change) between patients with ‘normal’ (<0.10) and ‘abnormal’ (>0.10) serum IL-1 levels. The relative abundances were estimated by matching the metagenomic sequence to the SEED gene function database using MG-RAST. The left-hand side indicates gene categories significantly more abundant in individuals with ‘abnormal’ serum IL-1 levels compared with ‘normal’, while the right side indicates the reverse.

Particularly noteworthy was the higher relative abundance of genes associated with nitrosative stress enzymes. These genes are directly involved in the reduction of Nitric Oxide (NO) free radicals. NO is used as a toxic defense against infectious organisms and regulates the growth and activity of inflammatory cells such as macrophages (35). Nitric oxide is involved in host defense and nitrosative stress has been established as a biomarker in the inflammatory and immune response (36,37). Studies have shown the biofilm in the periodontal disease has the capability to convert nitrate in the oral cavity to nitrite, which in turn is denitrified to nitric oxide (36). A 2009 study of 60 periodontitis subjects measured the NO levels in serum and found elevated levels of nitrite in periodontitis subjects compared to healthy subjects; Menaka *et al.* further described that NO presence reflects bone resorption leading to disease progression (36). Thus, the higher relative abundance of nitrosative stress enzymes in the biofilm of patients with higher IL-1ß is consistent with the research showing *F. nucleatum* stimulates IL-1ß and NO production.

## Conclusions

The results of our metagenomic and vascular function study provide evidence of a relationship in young adults between periodontal disease and specific measures of vascular dysfunction. The connection with specific members of the pathogenic oral community, particularly *Fusobacterium nucleatum*, corresponds well with previous human and mouse model studies and suggests a possible causal relationship between periodontitis and vascular dysfunction predisposing to atherosclerotic heart disease. While the connections appear robust, the sample size of our study is relatively modest. With the decrease in costs associated with high-throughput sequencing and cytokine analyses, much larger studies will be possible. Also, a study over a longer timeframe should allow greater opportunity for resolution of periodontitis and subsequent changes in measures of CVD. In the future, larger studies should be carried out with a full panel of metagenomics, serum and gingival crevicular cytokine analyses and CVD measurements over a longer time frame. These findings establish a clear relationship between the periodontal pathogen *F. nucleatum* and IL-1β, which has been shown to contribute significantly to heart disease. This study was conducted in adults under age 40 and suggests that decades of exposure to inflammatory cytokines contributes to life-threatening heart disease. Further study is needed to show that heart disease progression can be delayed with appropriate oral hygiene.

## Acknowledgements

We would like to thank the staff at the American Indian/Alaska Native (AIAN) population clinic, specifically to the medical, dental and administrative staff for their hard work and dedication. We also thank Rosalin Le for help coordinating sample collections and performing LDF measurements. We also thank the many willing participants in this study. This study was funded by a grant from the National Institutes of Health.

## Supporting Information

**S1 File. Data and analysis files**. Compressed collection of data files and analysis scripts used to generate the results of the study.

